# Gestational inhibition of CSF1R signaling using PLX5622 drives musculoskeletal changes in postnatal offspring

**DOI:** 10.64898/2025.12.16.694699

**Authors:** Rouzbeh Ostadsharif Memar, Matthew Rosin, Siddharth R. Vora, Jessica M. Rosin

## Abstract

Despite our understanding of the musculoskeletal system under homeostatic conditions and during tissue remodeling, the interplay between muscle and bone development in response to gestational perturbations is less well understood. Here, we used the colony-stimulating factor-1 receptor (CSF1R) inhibitor PLX5622 to disrupt macrophage and osteoclast proliferation, differentiation, and survival during the embryonic period in order to study the impacts on craniofacial development using high-resolution microcomputed tomography (μCT). Cranioskeletal and mandibular size and shape were assessed using geometric morphometric (GM) analysis and dense correspondence analysis (DeCA), while contrast-enhanced μCT and DeCA were utilized to examine the consequences of prenatal CSF1R inhibition on P1 offspring musculature. Combined, we observed significant disruptions to cranioskeletal and mandibular morphologies, and notable changes in the shape of the muscles of mastication and tongue in newly born pups exposed to the CSF1R inhibitor PLX5622 *in utero*. By assessing the craniofacial skeleton and associated musculature at birth, we provide a more direct view of how inhibition of CSF1R signaling across embryogenesis contributes to changes in musculoskeletal development during periods of craniofacial morphogenesis.

## INTRODUCTION

Colony-stimulating factor 1 receptor (CSF1R) is a receptor tyrosine kinase that directs the proliferation, differentiation, and survival of the mononuclear phagocyte lineages, including macrophages and osteoclasts, upon binding its ligands CSF1 and IL-34 (1,2). CSF1R signaling regulates organogenesis and homeostasis in multiple systems, ranging from microglial control of brain development to macrophage-mediated remodeling in the mammary gland, intestine, pancreas, and reproductive organs (3–6). Within the musculoskeletal system, CSF1R signaling governs osteoclast differentiation and bone remodeling, coordinates osteoblast–osteoclast coupling, and contributes to muscle development and repair by supporting macrophage-dependent myogenesis (7–10). Given these diverse functions, disrupting CSF1R signaling is predicted to impact craniofacial morphogenesis, where skeletal and muscular tissues develop in tight coordination. Indeed, genetic knockout animal models provide evidence for these roles. In mice, homozygous loss-of-function mutations in *Csf1* (*Csf1^op/op^*) or *Csf1r* (*Csf1r^−/−^*) eliminates or severely reduces macrophages and osteoclasts, leading to generalized osteopetrosis, delayed tooth eruption, abnormal bone matrix deposition, and frequent fractures (11–15). Craniofacial anomalies in these models include doming of the cranial vault, defective sutural biology, and disorganized intramembranous and endochondral ossification (14,15). Rats with the osteopetrotic mutation *toothless* (*tl*) or *Csf1r* knockout show similar sclerotic bone phenotypes, impaired ossification centers, and under-modelled cranial bases (9,16,17). Beyond skeletal consequences, CSF1R deficiency reduces muscle-resident macrophages and is associated with diminished muscle mass and delayed postnatal myofiber growth, underscoring the dual skeletal and muscular impact of the pathway during development (9,18,19).

Although genetic models highlight the indispensability of CSF1R, they carry limitations such as perinatal lethality, and permanent loss of signaling precludes stage-specific analyses (20). To overcome these challenges, small-molecule inhibitors such as PLX5622 have been used to achieve temporally controlled and reversible blockade of CSF1R (21,22). Pharmacological inhibition of CSF1R in mice during embryogenesis using PLX5622 has revealed craniofacial abnormalities at and shortly after weaning at postnatal days 21 (P21) and P28, including doming of the cranial vault, notched incisors with ectopic enamel ridges, and altered molar cusp morphology (21,23). However, assessment of craniofacial phenotypes at these endpoints does not resolve whether the abnormalities originate directly from embryonic disruptions or instead reflect cumulative postnatal adaptations. Moreover, postnatal musculature was not assessed following embryonic CSF1R inhibition using PLX5622. To address these gaps in our knowledge, we employed a pharmacological strategy to inhibit CSF1R signaling during embryogenesis using PLX5622 and characterized craniofacial phenotypes one day after birth at P1. Here, we comprehensively investigate the impact of prenatal CSF1R inhibition on cranioskeletal and mandibular size and shape using high-resolution microcomputed tomography (μCT), geometric morphometric (GM) analyses, and dense correspondence analysis (DeCA). We also use contrast-enhanced μCT and DeCA to examine the impact of prenatal CSF1R inhibition on the key facial musculature of P1 offspring, including the muscles of mastication and tongue. Herein, we demonstrate that CSF1R inhibition *in utero* significantly disrupts cranioskeletal and mandibular morphologies in newly born pups, with pronounced changes in the shape of the musculature also observable in P1 PLX5622 offspring. Through our assessment of the craniofacial skeleton and associated soft tissues at birth, we help clarify how inhibition of CSF1R signaling during the embryonic period contributes to craniofacial development, offering new insights into the basis of the craniofacial anomalies seen in both animal models and human CSF1R-related disorders such as brain abnormalities, neurodegeneration, and dysosteosclerosis (BANDDOS) (24–27).

## RESULTS

### Gestational inhibition of CSF1R signaling alters the shape of the cranial skeleton and mandible

As gestational exposure (embryonic day 3.5 (E3.5) to birth) to the CSF1R inhibitor PLX5622 reduces craniofacial macrophage populations to <50% and disrupts osteoclastogenesis, resulting in widespread craniofacial disruptions impacting embryonic bone development (28), we were interested in studying the shape of both muscles and bones in P1 offspring in response to prenatal CSF1R inhibition. To avoid artifacts caused by postmortem jaw position, cranial and mandibular shapes were analyzed separately using geometric morphometrics (GM). The centroid size of the cranial skeleton did not differ significantly between P1 CD1 control and PLX5622 diet mice (Fig. 1A; p=0.144). However, principal component analysis (PCA) identified a strong association between diet (control vs. PLX5622) and skull shape. Indeed, PC1, which explained 27% of total variation, was significantly correlated with diet (Fig. 1B), and canonical variate analysis further demonstrated broad segregation between control and PLX5622 diet animals (Fig. 1C). Wireframes depicting the shape changes associated with PC1 highlighted doming of the posterior cranial vault as a defining phenotype in P1 CD1 PLX5622 pups (Fig. 1D). Moreover, multivariate analysis of variance (MANOVA) confirmed that the first six PCs, together accounting for approximately 80% of the variation, were significantly correlated with diet (p=0.013; Table 1). Notably, no PCs were significantly associated with sex, and MANOVA did not reveal a sex-dependent effect (data not shown).

**Figure 1.**
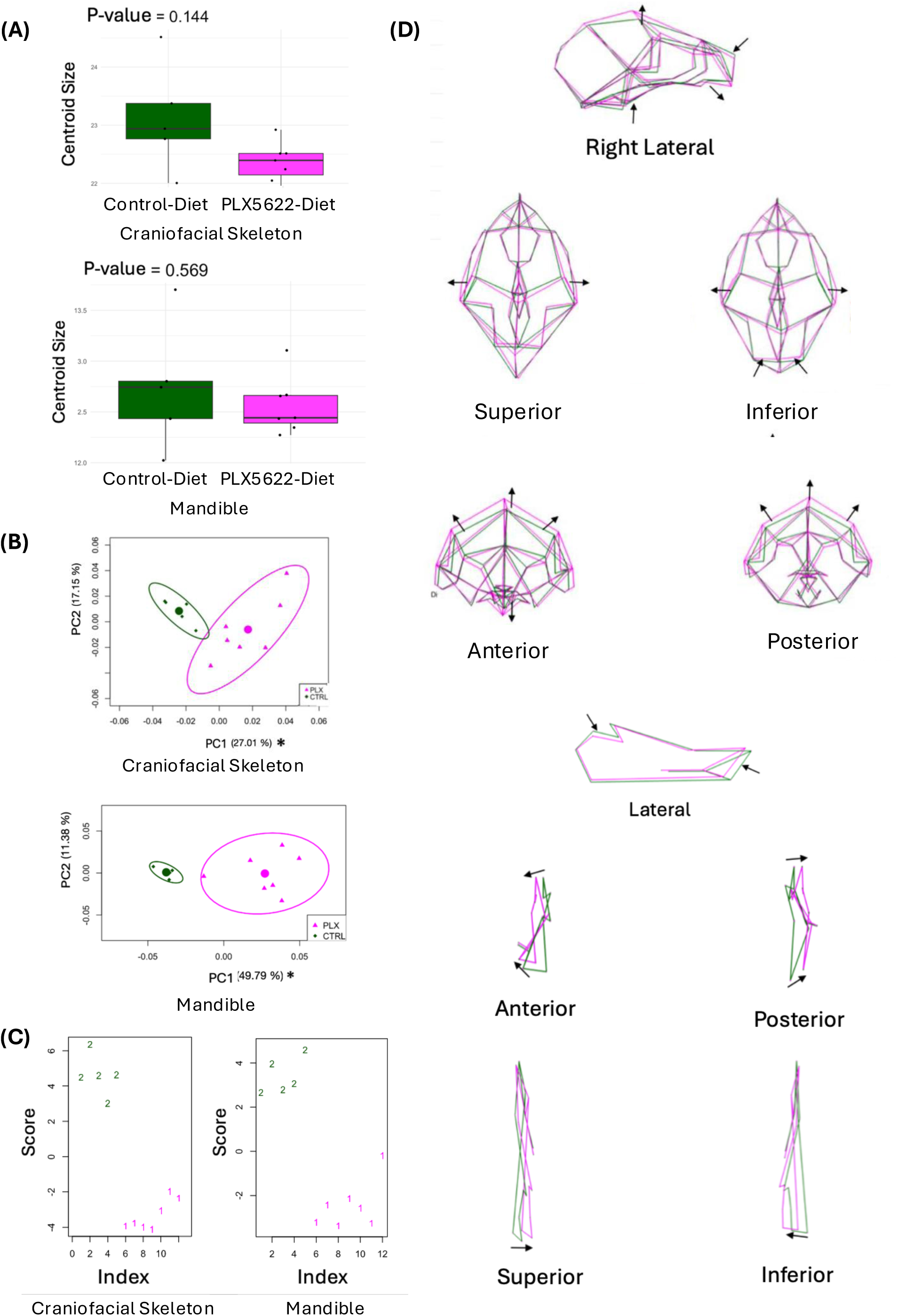
GM analysis for the cranial skeleton and mandible of P1 CD1 control and PLX5622 diet offspring. (A) Box plots comparing the centroid sizes of the cranial skeleton landmarks (top) and mandible landmarks (bottom) for P1 control and PLX5622 diet pups (green and pink bars, respectively), with the means denoted by a dark black line. (B) PCA plots of the cranial skeleton landmarks (top) and mandible landmarks (bottom) for P1 CD1 control and PLX5622 diet pups. PC1 and PC2 scores are plotted along the x-and y-axis, respectively. Control (CTRL) and PLX5622 (PLX) animals are represented by green diamonds and pink triangles, respectively. Green and pink circles represent the average shape configuration for each group, with 90% confidence ellipses depicted in the corresponding color for each group. The PC that significantly correlated with diet is marked by an asterisk. (C) Plots showing the distribution of canonical variate scores, with control pups depicted in green ‘2’ and PLX5622 pups depicted as pink ‘1’ values, for the cranial skeleton (left) and mandible (right). (D) Wireframes depicting the shape variation along the first PC for the cranial skeleton (top) and mandible (bottom), with green depicting control diet pups and pink depicting PLX5622 diet pups, displayed from different angles. Arrows depict the direction of shape change in the PLX5622 diet wireframe compared to the control diet wireframe along the first PC.

**Table 1.** MANOVA results for the cranial skeleton and mandible shape variation between P1 CD1 control and PLX5622 diet pups. MANOVA for cranial skeleton landmark coordinates (top) and mandible landmark coordinates (bottom), using the loadings from the first 6 PCs of shape variation between mice when testing the variable diet (CTRL vs PLX5622) using both the Pillai and Hotelling-Lawley statistics.

Craniofacial Skeleton
| Factor | Df | Pillai | Hotelling–<br>Lawley | approx. F | num Df | den Df | Pr(>F) |
| --- | --- | --- | --- | --- | --- | --- | --- |
| Diet | 1 | 0.91942 | 11.411 | 9.509 | 6 | 5 | 0.013 |
| Residuals | 10 | — | — | — | — | — | — |

Mandible
| Factor | Df | Pillai | Hotelling–<br>Lawley | approx. F | num Df | den Df | Pr(>F) |
| --- | --- | --- | --- | --- | --- | --- | --- |
| Diet | 1 | 0.86582 | 6.4524 | 5.377 | 6 | 5 | 0.042 |
| Residuals | 10 | — | — | — | — | — | — |

Interesting, a similar pattern was observed in the mandible. Centroid size was similar for the P1 CD1 control and PLX5622 diet mice (Fig. 1A; p=0.569), while PCA identified a significant association between diet and mandible shape (Fig. 1B). Specifically, PC1, explaining ∼50% of the overall variation in mandibular shape across specimens, was significantly correlated with diet (Fig. 1B), and canonical variate analysis showed complete segregation between P1 CD1 control and PLX5622 diet animals (Fig. 1C). Wireframes depict right hemi-mandible shape changes associated with PC1, highlighting an underdevelopment of the mandible in P1 CD1 PLX5622 pups (Fig. 1D). MANOVA confirmed that the first six PCs, representing ∼80% of variation, were significantly associated with diet (p=0.042; Table 1), while sex had no significant effect on mandibular shape (data not shown).

To further localize these GM-identified shape differences across the full bone surface, we applied Dense Correspondence Analysis (DeCA) to map surface-level shape variation between P1 control and PLX5622-exposed pups. DeCA of the cranial skeleton revealed discrete regions of deviation in PLX5622 animals, with the most prominent differences occurring along the superior and posterior cranial vault, consistent with the doming phenotype identified by GM analyses (Fig. 2A). Additional localized differences were observed in the otic region, with smaller differences visible in the anterior naso-maxillary complex and the cranial base (Fig. 2A). DeCA of the mandible similarly demonstrated regional shape differences between diet groups, with the greatest deviations occurring at the anterior mandibular tip, the lower border of the mandibular body, and the posterior ramal region (Fig. 2B). The later locations correspond closely with major muscle attachment sites, suggesting that early deficits in osteoclast-mediated remodeling at mechanically responsive regions may contribute to the mandibular phenotype at P1.

**Figure 2.**
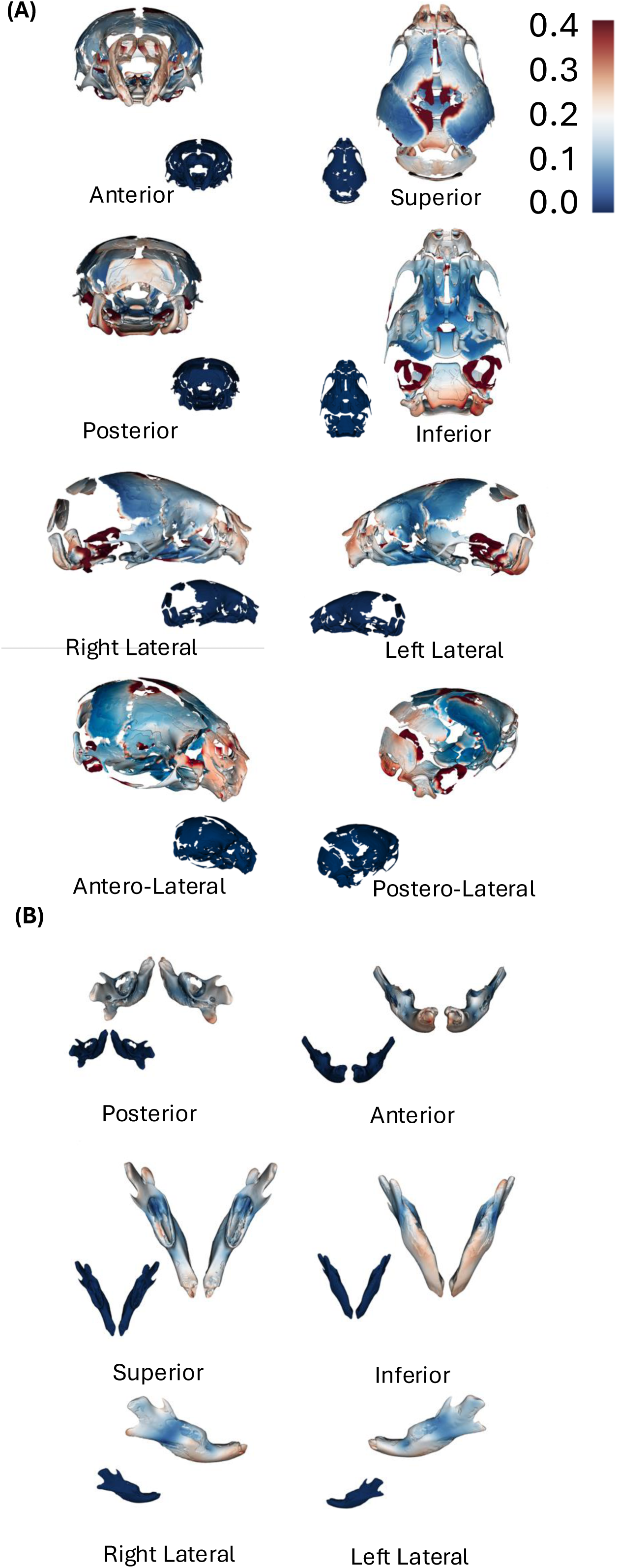
DeCA for the cranial skeleton and mandible of P1 CD1 control and PLX5622 diet offspring. (A-B) Images depict the mean cranial skeleton and mandible models for PLX5622 diet specimens compared to the mean control diet cranial skeleton and mandible models, which was used as the reference. The smaller insets in each of the panels represent the mean control model compared against itself. The bar on the top right of the diagram represents the magnitude of shape difference with its corresponding color. (A) Anterior view, posterior view, superior view, inferior view, right lateral view, left lateral view, antero-lateral view, postero-lateral view of the cranial skeleton are depicted. (B) Posterior view, anterior view, superior view, inferior view, right lateral view, and left lateral view of the mandible are depicted.

### Embryonic CSF1R inhibition alters the shape of the muscles of mastication and tongue

To study the impact of prenatal CSF1R inhibition on soft tissue morphology, we examined contrast-enhanced μCT reconstructions of the right-side muscles of mastication and tongue, and quantified surface level differences using DeCA. When analyzed individually (Fig. 3A), each structure displayed distinct regions of shape variation between groups.

**Figure 3.**
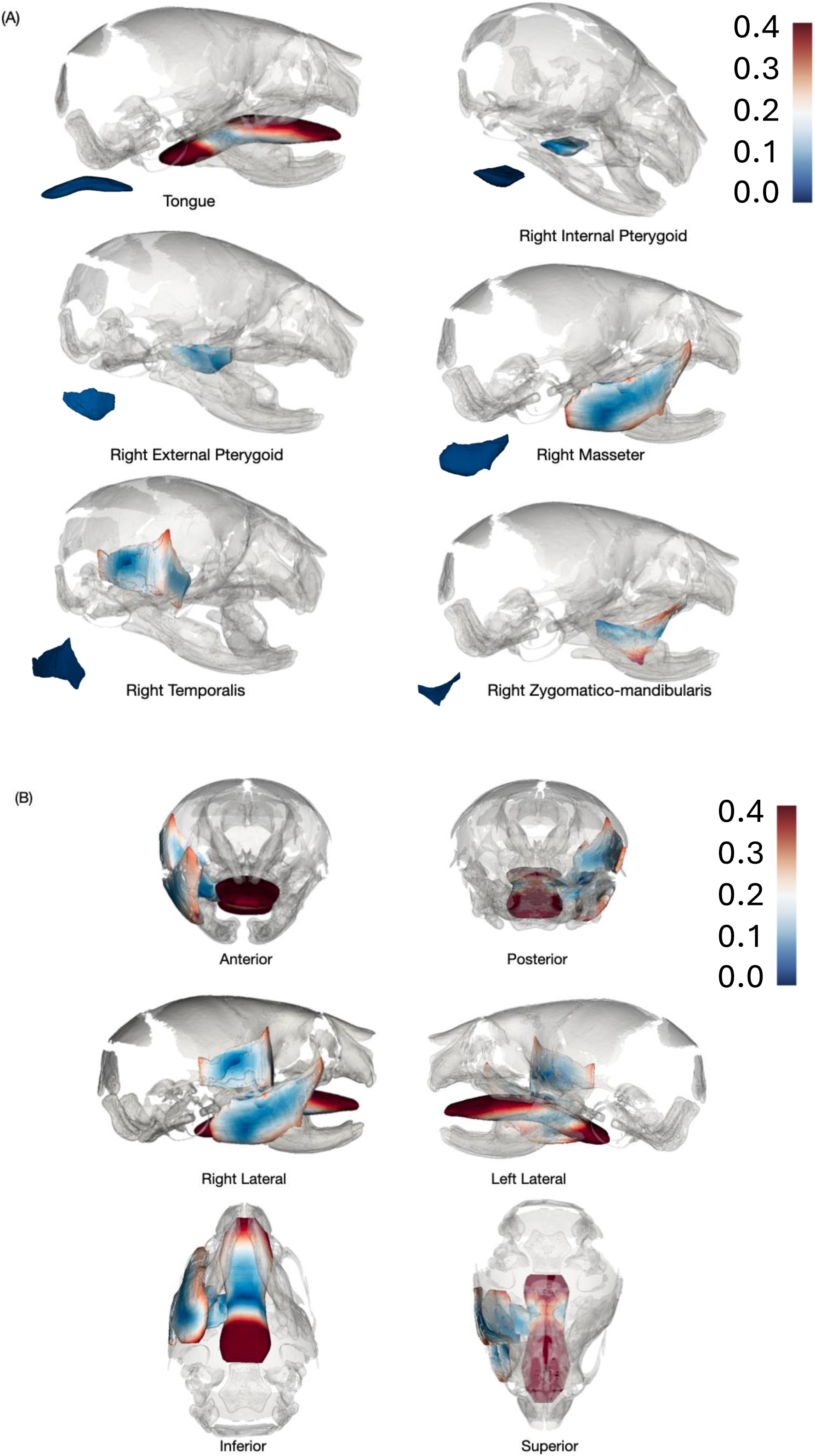
DeCA for the muscles of mastication and tongue of P1 CD1 control and PLX5622 diet offspring. (A-B) Images depict the shape changes of mean muscles of mastication and the tongue in PLX5622 diet specimens compared to control diet specimens, which were used as the reference. Bars on the right of the diagram represent the magnitude of shape difference with their corresponding color. (A) Representative images of shape changes for each individual muscle of mastication and tongue for PLX5622 diet specimens compared to control diet specimens, where the smaller insets in each of the panels represent the mean control model compared against itself. (B) Muscles of mastication and tongue models placed next to each other, representing the shape changes in PLX5622 diet specimens against the control diet specimens.

The masseter displayed pronounced shape differences along the superior border, where it inserts into the zygomatic arch and along the inferior and posterior borders at mandibular attachment sites (Fig. 3A). The zygomatico-mandibularis showed shape differences along the superior margin near the zygomatic arch and the inferior margin along the lateral aspect of the mandible (Fig. 3A). The temporalis exhibited shape variation along its posterior-superior border, corresponding to its cranial attachment site, and along its antero-inferior border where it attaches to the coronoid process of the mandible (Fig. 3A). The internal pterygoid displayed variation in shape along its medial border, at the cranial base and its lateral border where it inserts on the mandible, while the external pterygoid showed distinct shape differences along the anterior border near the alispheno-squamosal suture and posterior border adjacent to the tympanic bulla (Fig. 3A). In addition to the muscles of mastication, the tongue displayed notable shape differences between P1 CD1 control and PLX5622 pups across the anterior two-thirds of the dorsal surface and at both the dorsal and ventral regions of the base (Fig. 3A).

When the soft tissues are viewed on a common scale, the tongue exhibited the greatest magnitude of shape variation, whereas the pterygoid muscles demonstrated the least variation in shape between P1 CD1 control and PLX5622 pups (Fig. 3B).

### DeCA analysis identifies overlapping regions of skeletal and muscular shape variation following prenatal CSF1R inhibition

To compare skeletal and soft tissue surface variation, we performed DeCA overlays of each mean muscle of mastication and the tongue against the skull model, mapping regions of shape variation for both tissues using distinct color schemes (Fig. 4). This visualization enables spatial comparison of skeletal and muscular surface differences in P1 CD1 PLX5622 diet pups relative to controls. The masseter showed variation along its superior and inferior borders, corresponding to changes in the zygomatic arch and mandibular surface (Fig. 4). The temporalis exhibited shape differences along its anterior and posterior borders, which align with regions of variation in the lateral cranial vault (Fig. 4). Similarly, the zygomatico-mandibularis and pterygoid muscles displayed differences at both mandibular and cranial base interfaces, regions that were also highlighted in the skull model (Fig. 4). The tongue demonstrated deviations at its base and anterior dorsal surface that aligned with areas of variation in the anterior maxilla and mandible (Fig. 4).

**Figure 4.**
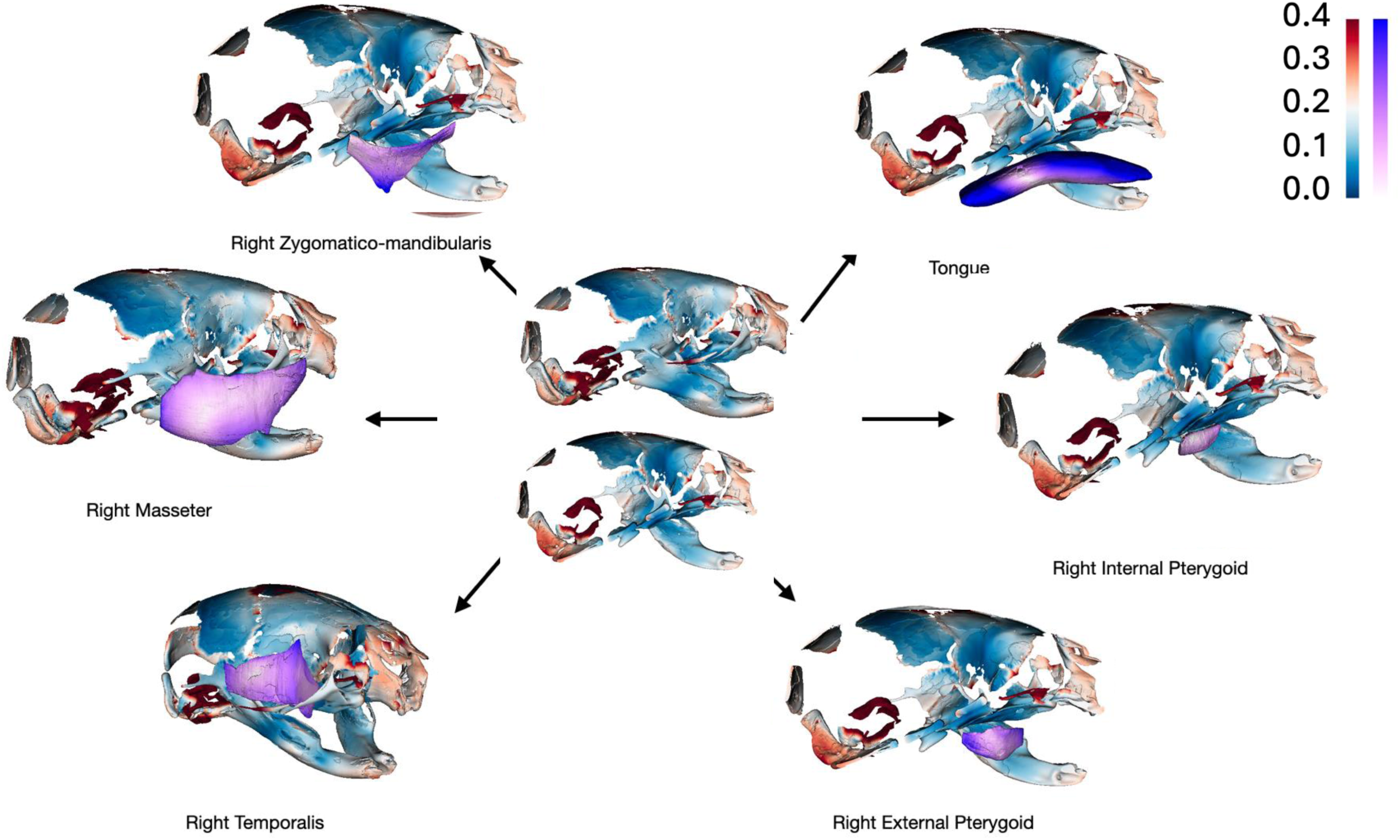
DeCA highlights shape changes for each muscle of mastication and tongue compared against shape changes identified for the cranial skeleton and mandible of P1 CD1 control and PLX5622 diet offspring. Images depict the mean muscles of mastication and tongue models for PLX5622 diet specimens compared to control diet specimens, placed individually against a complete head model or a sagittally sectioned model (for better visibility) representing the shape changes in mean PLX5622 diet head specimens compared to the mean control diet head specimens. The bars on the top right of the diagram represent the magnitude of shape difference with their corresponding colors.

## DISCUSSION

In this study, we demonstrate that prenatal inhibition of CSF1R signaling using PLX5622 alters the morphology of both craniofacial bones and associated soft tissues at birth. Using DeCA, we identified surface-level shape differences in the skull, mandible, muscles of mastication, and tongue. This methodology complements traditional GM approaches, which characterize shape differences using homologous landmarks, and overlooks surface features across regions lacking discrete anatomical points. This is particularly important for soft tissues such as the musculature as well as bony areas corresponding to soft tissue attachment sites. Muscles are pliable and continuously change shape in response to functional loading and attachment geometry; therefore, DeCA’s high-density maps provide a more physiologically relevant way to detect such alterations (29). Applying this technique allowed us to examine changes in both bone and muscle that would otherwise be difficult to characterize, establishing DeCA as a powerful tool for craniofacial phenotyping.

Our findings at P1 can be interpreted alongside our prior results at P21 given that the same pharmacological CSF1R inhibition protocol was employed across embryogenesis using PLX5622 (23,30). At both P1 and P21, a consistent phenotype emerged: a dome-shaped cranial vault and underdevelopment of the mandible. These overlapping features suggest that the cranial vault and mandibular phenotypes observed at P21 (23,30) likely arise from CSF1R inhibition embryonically. However, some differences exist between P1 and P21. Indeed, at P21 a significant reduction in centroid size of the craniofacial skeleton and mandible was documented (23), whereas no such size differences were detected at P1 in the current study. This pattern could suggest that prenatal CSF1R inhibition establishes an early defect in cranioskeletal shape and patterning at birth, while postnatal bone growth and remodeling in a compromised system amplify the phenotype into a measurable size difference by P21. Skeletal staining of P1 CD1 control and PLX5622 pups with Alizarin Red and Alcian Blue showed significant linear reductions in both skull and mandibular size (28). This suggests that differences in quantification methods (e.g., centroid-based versus linear measurements) may influence the observed size outcomes and should be interpreted with consideration of each method’s limitations. Importantly, PLX5622 exposure was restricted to embryogenesis, and CSF1R signaling resumes postnatally once the drug is withdrawn (31). The observed cranioskeletal shape phenotype at P1, and its amplification into measurable size deficits by P21, indicates that early CSF1R-dependent events establish a developmental trajectory that is not reversed when signaling is restored. These findings support a model in which embryonic disruption of osteoclast- and macrophage-mediated patterning creates an altered template upon which postnatal growth proceeds, thereby perpetuating and enlarging the phenotype rather than correcting it.

Interestingly, shape variation in the muscles of mastication and tongue consistently corresponded to their origins and insertions on the skull. For example, mandibular and cranial vault differences were mirrored in the temporalis and masseter, and anterior mandibular and premaxillary differences were paralleled by tongue shape variation. This parallel mapping underscores the tight developmental and functional integration between hard and soft tissues during craniofacial morphogenesis. It also illustrates how changes in one system can cascade into the other: altered bone geometry may impact attachment sites, which can change muscular morphology, while altered muscular growth and force transmission may, in turn, influence bone growth trajectories (32). Based on the collective results from this study and our prior analysis (21,23,28), we propose that the skeletal changes likely precede muscular alterations. Indeed, CSF1R signaling is critical for osteoclast differentiation and function, and osteoclasts, in turn, regulate osteoblast differentiation and bone matrix deposition (7,11–15). Disruption of this signaling axis during embryogenesis impairs ossification and sutural development (28), resulting in the domed cranial vault and underdeveloped mandible observed in this study. The muscular alterations are likely secondary to the underlying skeletal defects. Previous work has shown that embryonic inhibition of CSF1R signaling leads to defects in *Wnt1+* neural crest derivatives, Sp7+ osteoblasts, and mineralization within the premaxilla and mandible of embryonic day 15.5 (E15.5) embryos, whereas no significant differences in muscle development were detected between E11.5 and E15.5 (28). It is important to note that muscle-resident macrophages are also dependent on CSF1R signaling and can play trophic roles in growth (e.g., via IGF-1 and related cues); therefore, loss of these macrophages could reduce muscle volume and maturation of the musculature later in embryogenesis (9,18,19).

Although the consistent mapping of muscular changes to skeletal attachment points suggests that altered bone geometry is a primary driver of the soft-tissue phenotypes, this study has several limitations. For example, hard and soft tissue morphology could not be evaluated within the same animal due to interference from PTA staining, preventing fully integrated analyses, and μCT resolution did not permit segmentation of smaller or layered muscles, potentially masking subtle group-specific differences. Moreover, DeCA, while powerful, provides measures of magnitude but not directionality of shape change, constraining interpretation of some heatmap results. Accordingly, longitudinal imaging approaches, such as serial μCT or MRI, could track how embryonic defects evolve into postnatal size and shape changes, and higher-resolution imaging combined with histological validation could clarify the cellular mechanisms underlying the changes detected by DeCA. It is also important to note that all experiments were conducted in CD1 mice and strain-dependent variability in craniofacial phenotypes resulting from CSF1R inhibition have been reported (11,14,28,33). Therefore, future work could perform a comparative study across multiple strains to help reveal how genetic background modulates the craniofacial consequences of CSF1R inhibition *in utero*.

By examining the impact of prenatal CSF1R inhibition on craniofacial morphogenesis, we demonstrated striking musculoskeletal shape changes in P1 CD1 PLX5622 pups. Additionally, we showed that cranioskeletal and muscle shape changes are closely mapped to one another, supporting a model in which skeletal defects arise first, and muscular abnormalities follow. Through our utilization of DeCA, we established a novel framework for analyzing both hard and soft tissues in parallel, where P1 emerges as a critical developmental time-point for understanding the origins of craniofacial phenotypes arising from CSF1R inhibition *in utero*, bridging embryonic disruptions with postnatal outcomes we and others have observed at later ages (2,8,9,11–14,16,23,28).

## AUTHOR CONTRIBUTIONS

S.R.V. and J.M.R. conceived and conceptualized the study. R.O.M., M.R. and J.M.R. generated the samples used in the study. R.O.M. and M.R. prepared the samples for μCT scanning. R.O.M. and S.R.V. performed all analysis. R.O.M. and J.M.R wrote the manuscript and S.R.V. contributed to data interpretation and manuscript editing. All authors approved the final manuscript.

## ACKNOWLEDGEMENTS

The authors thank the UBC Centre for High-Throughput Phenogenomics—a facility supported by the Canada Foundation for Innovation, British Columbia Knowledge Development Foundation, and the UBC Faculty of Dentistry—for help with µCT scans, and the UBC Centre for Disease Modeling for animal care. We would also like to acknowledge Felix Ma from the Rosin lab for help with sample preparation and critical discussion, and the Richman laboratory for providing PTA and for critical discussions. This work was supported by an NSERC Discovery Grant to J.M.R. (RGPIN-2022-03718). J.M.R. is a Michael Smith Health Research (MSHR) BC Scholar and a Tier 2 Canada Research Chair (CRC) in Immune Regulation of Developmental Programs.

## DATA AVAILABILITY

The data that support the findings presented in this study are available from the corresponding authors upon reasonable request.

## MATERIALS & METHODS

### Mouse handling and CSF1R inhibition

Animal handling and specimen processing were carried out in accordance with guidelines and regulations of the Canadian Council of Animal Care and received prior approval from the University of British Columbia’s Animal Care Committee (protocols A21-0170 and A21-0171). Pregnant CD1 dams (strain code 022, Charles River) were administered the Plexxikon CSF1R inhibitor PLX5622 (1200 PPM added to chow AIN-76A, Research Diets) starting at E3.5 and continuing until birth. Control CD1 dams received control diet (AIN-76A, Research Diets) for the same duration. P1 CD1 pups from control and PLX5622 diet dams were euthanized within 24 hours following birth via decapitation. The heads and bodies were frozen at −20 °C in 15 mL Falcon tubes. P1 CD1 control diet pups were generated from three separate litters and P1 CD1 PLX5622 diet pups were generated from two separate litters.

### Genotyping

DNA was isolated from the tail tissue collected from each P1 CD1 pup. Target genes were amplified using PCR, and the sex of each mouse pup was determined by assessing the size of amplified DNA fragments on an agarose gel. The sex genes assessed were *X-linked lymphocyte-regulated* (*Xlr*) and *Sycp3-like-Y-linked* (*Sly*). The size of the bands associated with amplification of these genes were: X chromosome, 685 bp (*Xlr*); Y chromosome, 280 bp (*Sly*) (34).

### Phosphotungstic acid staining

P1 CD1 pups were fixed overnight with 4% paraformaldehyde at 4 °C. The samples were then dehydrated through serial incubations with increasing methanol concentrations (30%, 50% and 70%) in PBS. Subsequently, each sample was incubated with 1% phosphotungstic acid (PTA) in a 90% methanol solution for two weeks, ending with rehydration through serial incubations with decreasing methanol concentrations (70%, 50% and 30%). Incubation in each of the methanol concentrations described above, at both the dehydration and rehydration steps, was done for two days (35). In total, 7 control diet P1 CD1 pups (3 males and 4 females) and 6 PLX5622 diet P1 CD1 pups (3 males and 3 females) were stained with PTA.

### Micro-CT scanning and image re-orientation

After embedding the heads in 1% agarose, the PTA-stained heads (7 control diet and 6 PLX5622 diet P1 CD1 pups) were scanned using micro-CT (μCT) at the Centre for High-Throughput Phenogenomics at the University of British Columbia using a Scanco Medical μCT100 scanner at 55 kVp and 200 μA with a resolution of 7 µm. The output was exported as DICOM files. Additionally, 5 control diet (3 males and 2 females) and 7 PLX5622 diet (4 males and 3 females) P1 CD1 heads were scanned without PTA-staining, at a resolution of 15 µm after they were thawed at room temperature (RT) for one hour and immobilized in agarose.

DICOM files output from the scans were first re-oriented for visualization, using the image transformation function in Amira software [Version 2022.2]. The re-orientation was performed in order to position all mouse pup heads along a similar anterior–posterior, medio-lateral, and superior–inferior axis, centered within the morphospace. The transformed images were re-sampled and exported as new DICOM stacks. All subsequent analyses were performed on the re-sampled DICOM files imported into 3D Slicer [Version 5.2.2].

### Data segmentation and landmarking

The manual and semi-automated segmentation tools based on thresholding in 3D Slicer 5.2.2 were used to segment the skulls of the non-PTA-stained specimens. To isolate true shape variation from postmortem shifts in mandibular position, we conducted separate geometric morphometric (GM) analyses on landmark configurations placed on the mandible and the cranial skeleton, which included the cranial vault, base, and midfacial regions (23) (Fig. S1). Similarly, dense correspondence analysis (DeCA; see below) was performed separately on the mandible and the cranial skeleton. Using the scans obtained from the PTA-stained specimen, muscles of mastication (masseter, temporalis, internal pterygoid, external pterygoid, zygomatico-temporalis) and the tongue were segmented on both the left and right sides of the head based on published literature (36).

A total of 65 landmarks were annotated on each skull model (45 on the cranial skeleton and 20 on the mandible; Fig. S1). Landmarks were also placed on all muscles of mastication, tongue, and the skull using 3D Slicer, on 3D models exported from segmentations (Fig. S2). Cranial and mandibular landmarks followed published definitions (37), which were limited to those which could be located reliably at P1. Since no comprehensive landmark sets exist in the literature for mouse muscles of mastication or tongue; landmarks were defined *de novo* based on reproducible, anatomically distinct features within segmentations. Seven landmarks were placed on the masseter, six on the temporalis, six on the zygomatico-mandibularis, four on the internal pterygoid, six on the external pterygoid, and ten on the tongue (Fig. S2).

Landmark placements were repeated after a 2-week interval to assess error. Euclidean distances between the two sets of coordinates were calculated across specimen. Mean and standard deviation values were calculated and any points exceeding one standard deviation above the mean were re-assessed to ensure correct landmark annotation (see Tables 2 and 3).

**Table 2.**
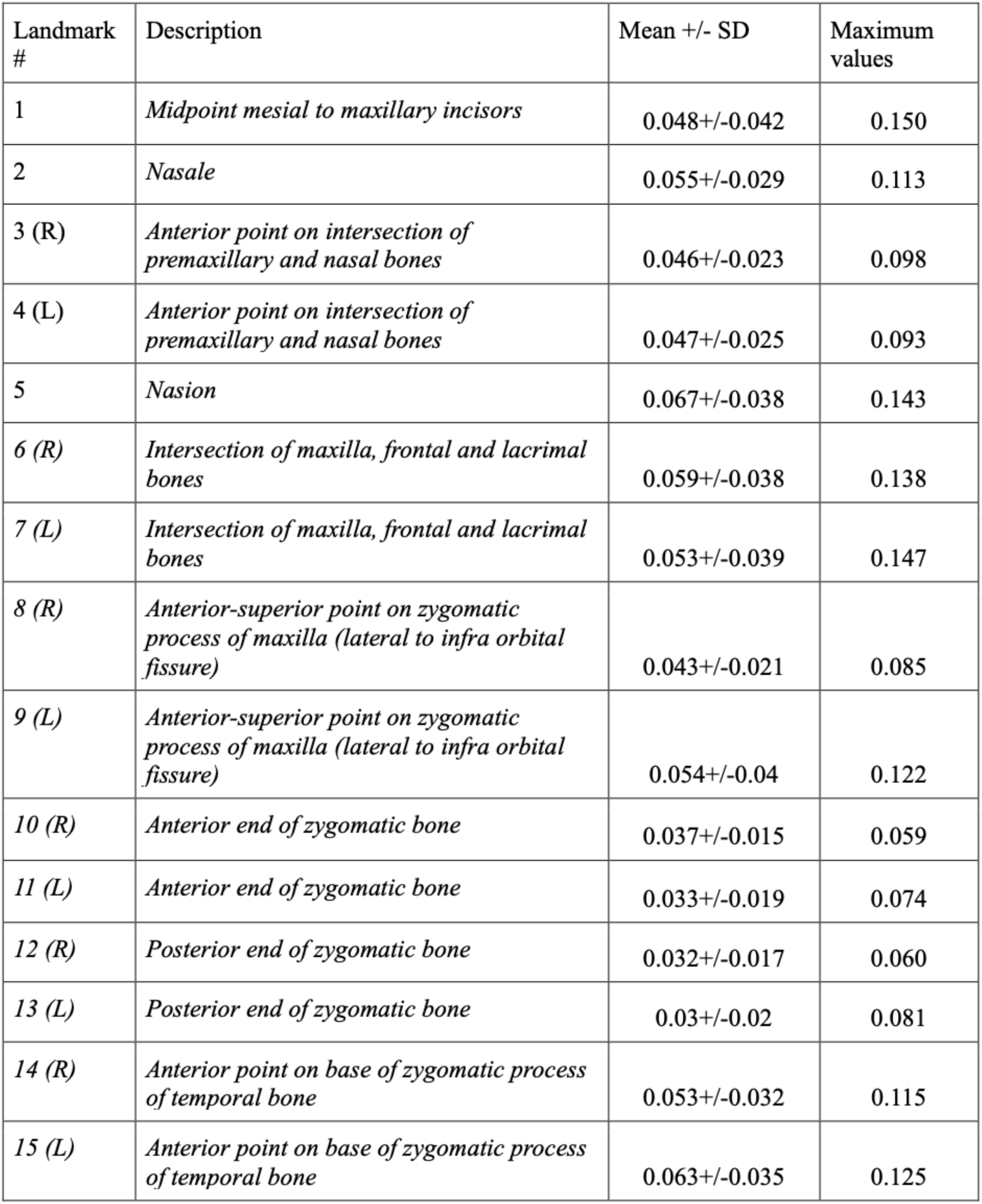
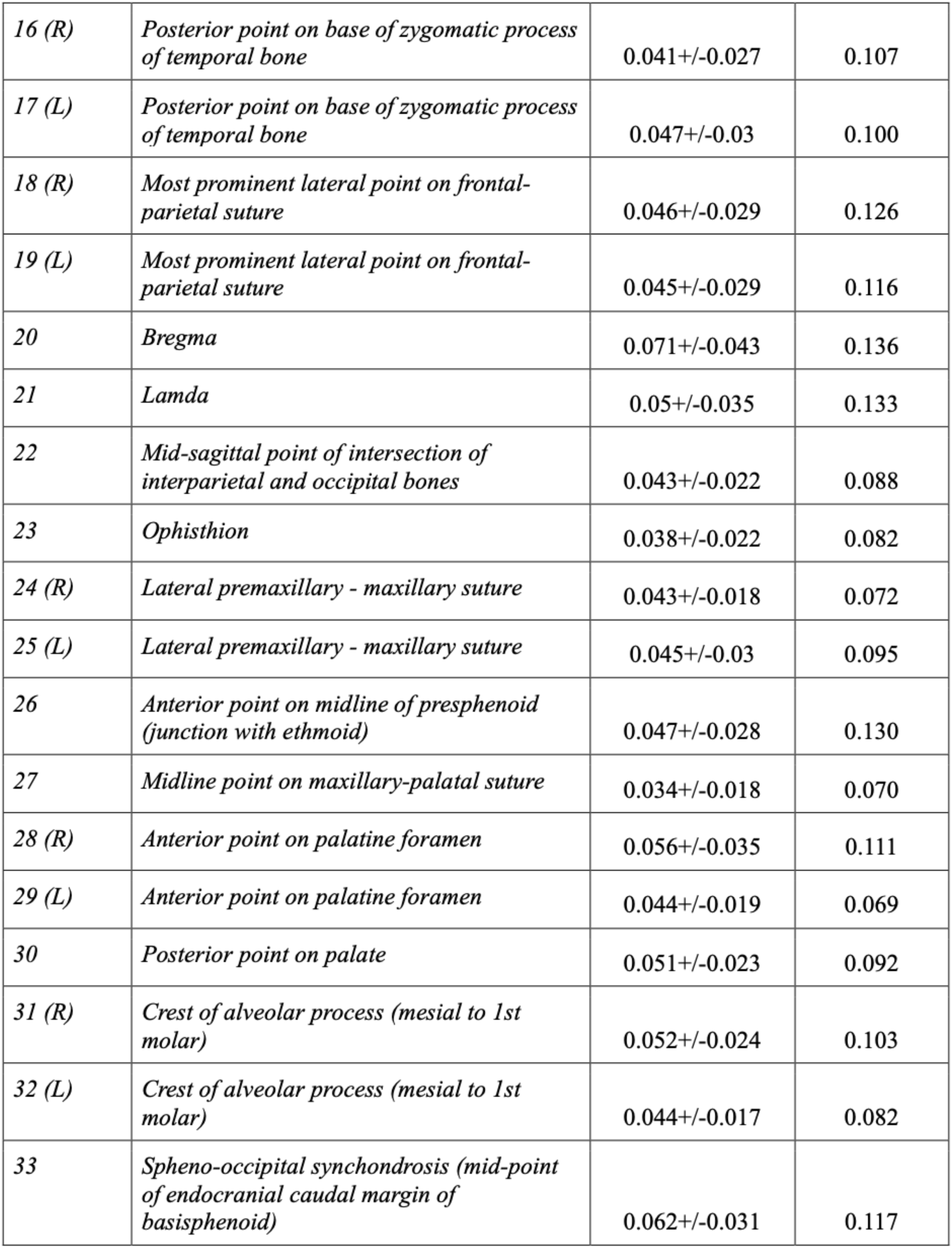
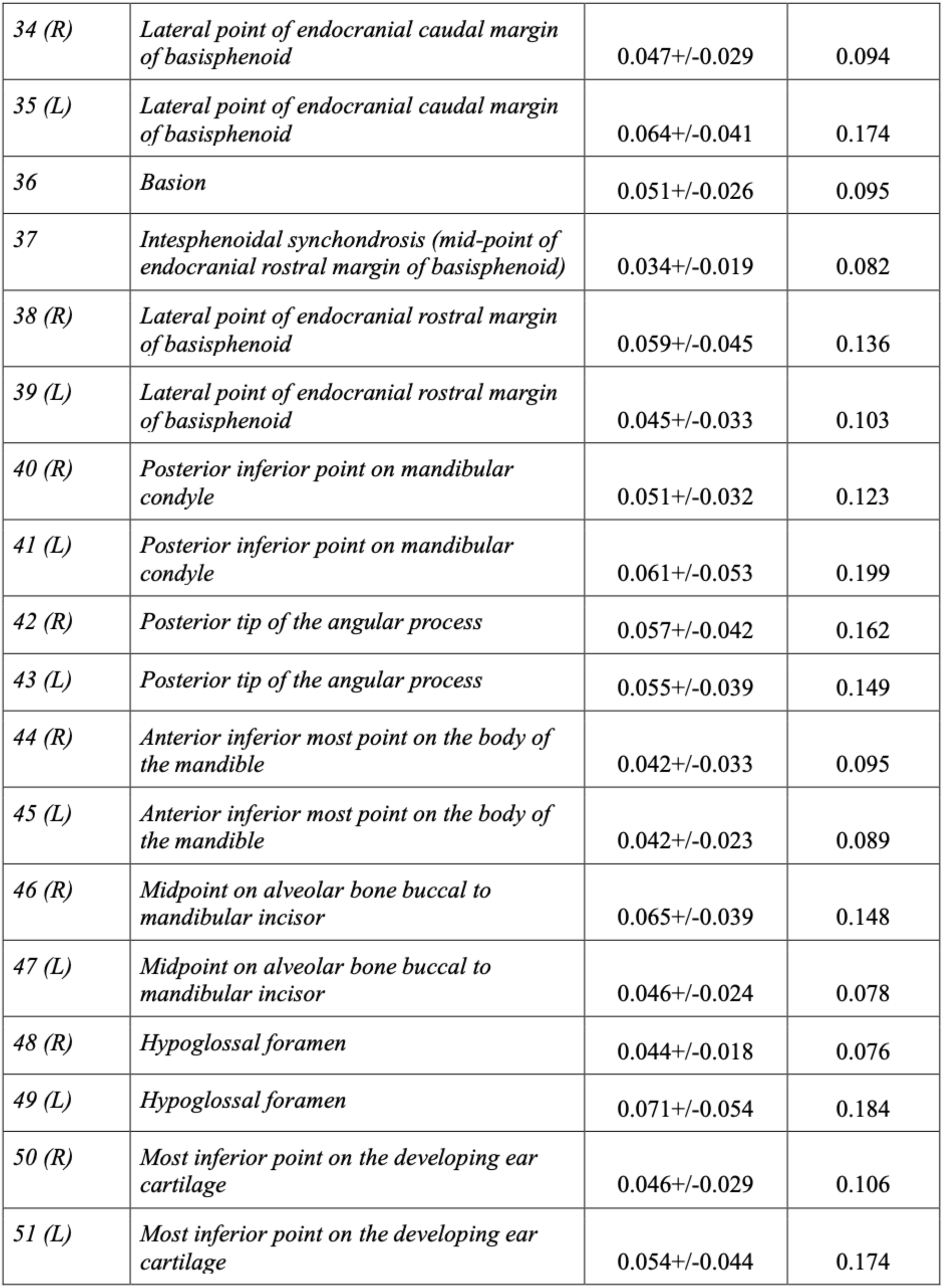
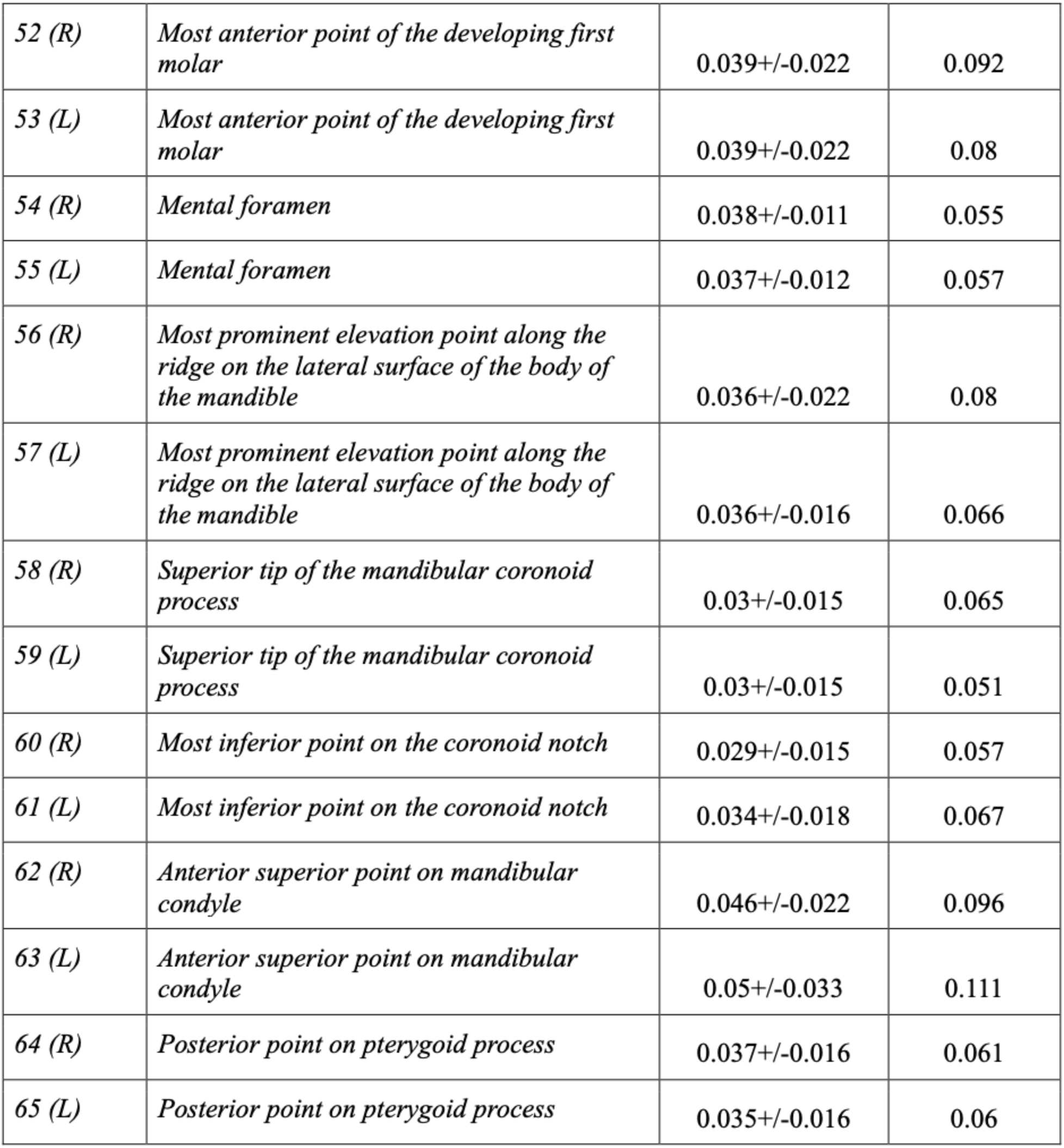
Description of anatomical landmarks placed on the skull along with their annotation error calculations. For the purposes of landmark annotation error assessment, the mean and standard deviation values for the distance between the first and second attempts at landmarking of all specimens are presented. The maximum value of the distance between the first and second attempts found for each landmark is also presented.

**Table 3.**
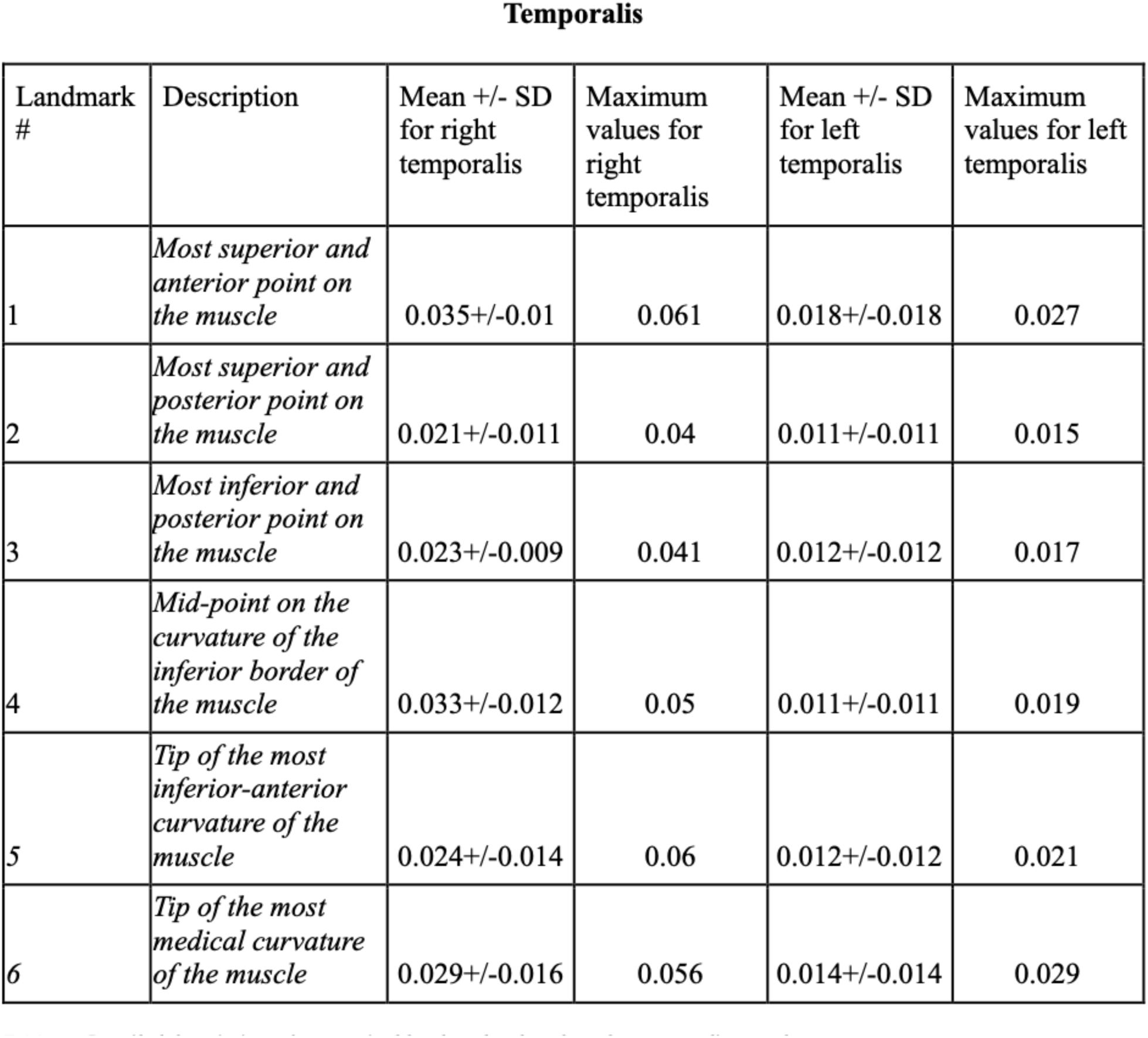
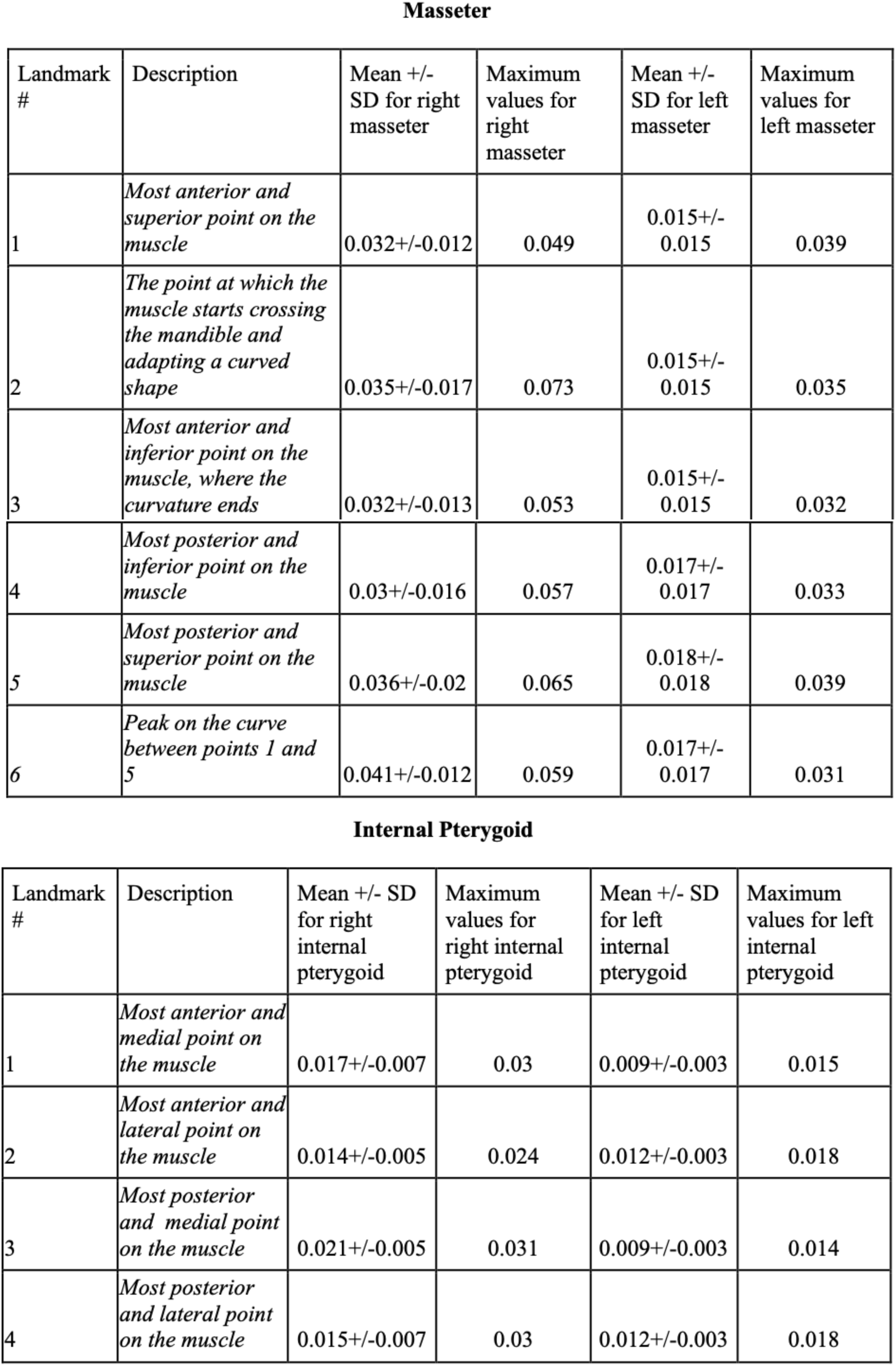
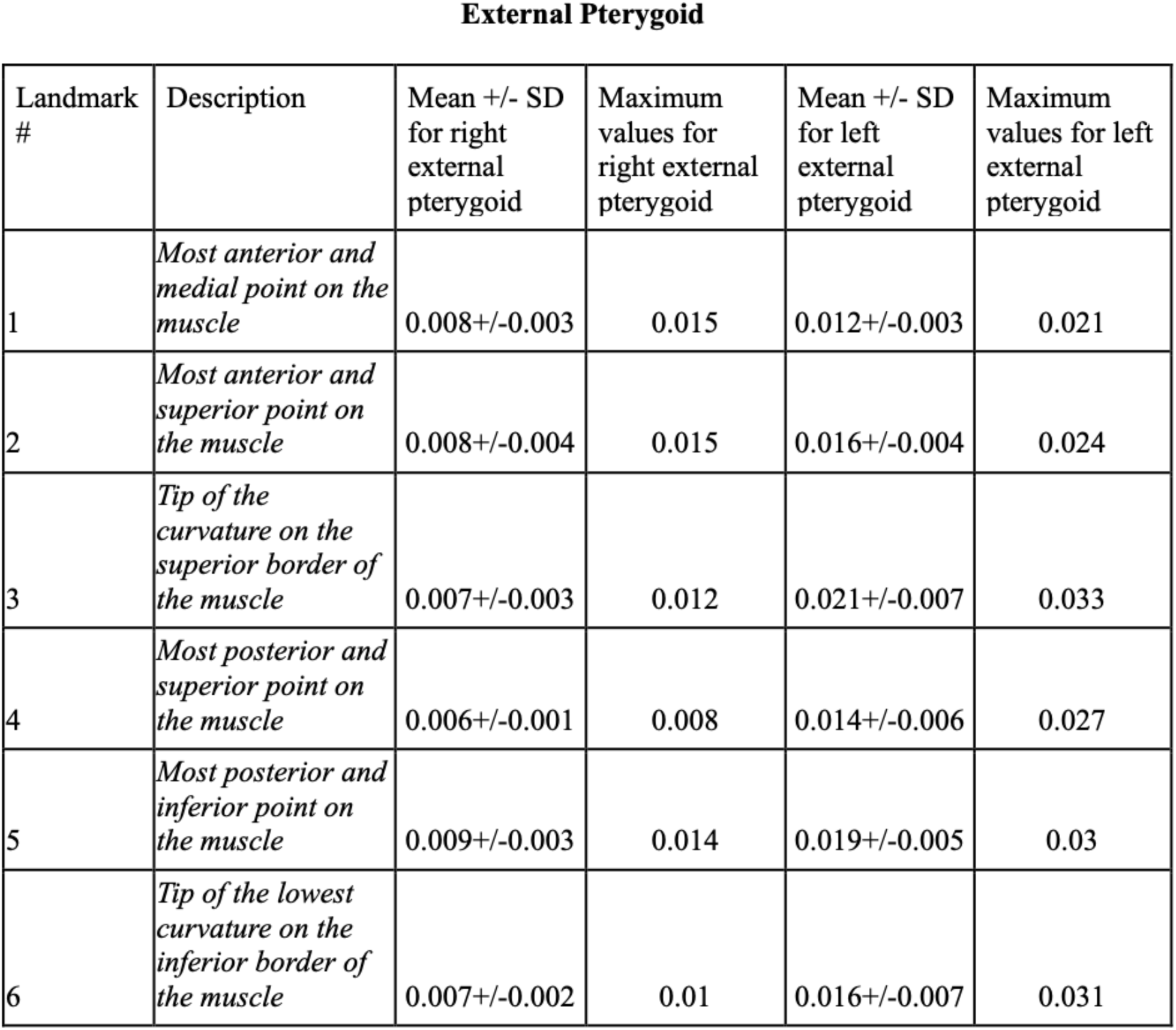
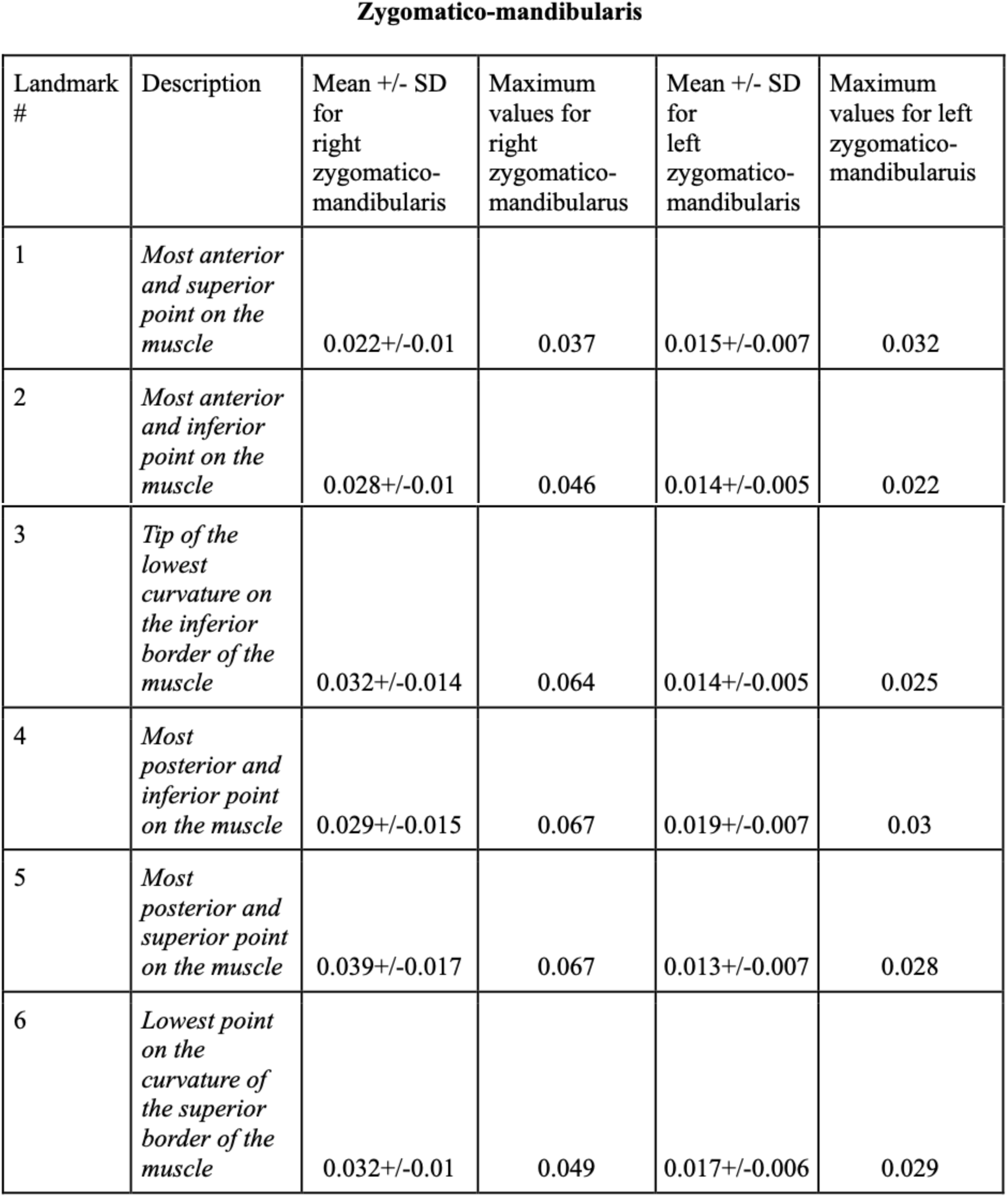
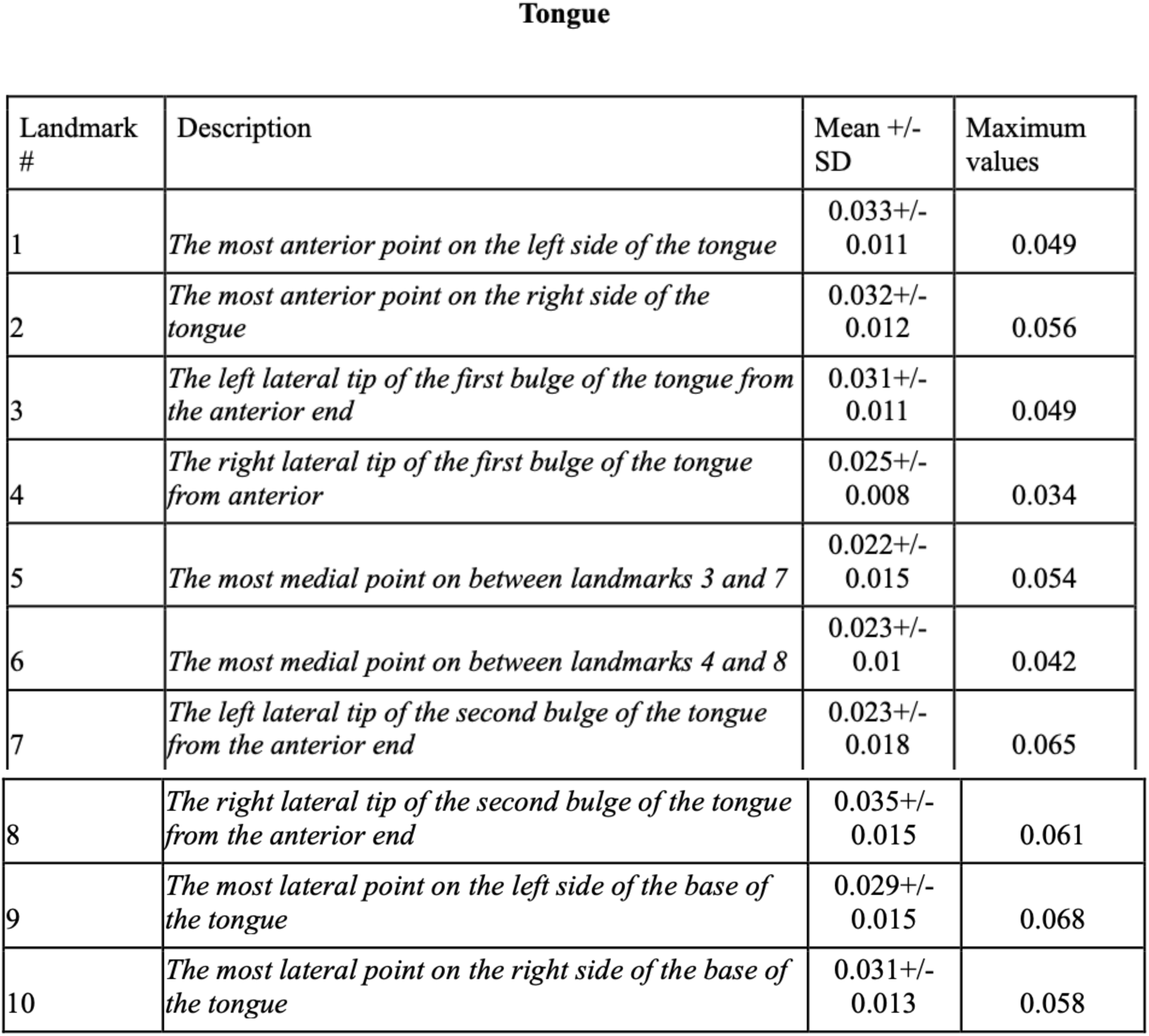
Description of anatomical landmarks placed on the muscles of mastication and tongue along with their annotation error calculations. For the purposes of landmark annotation error assessment, the mean and standard deviation values for the distance between the first and second attempts at landmarking of all specimens are presented. The maximum value of the distance between the first and second attempts found for each landmark is also presented.

### Geometric Morphometric Analysis

GM analysis was used to assess the shape differences in the cranial skeleton and mandible between control and PLX5622 diet specimens. Centroid sizes were calculated from landmark configurations and compared between diets using the Welch’s two-sample t-test (p<0.05; control diet vs. PLX5622 diet groups). Initial Procrustes superimposition was performed in order to correct for positional, rotational, and scaling of landmark configurations, yielding new Procrustes coordinates. Principal component analysis (PCA) was used to characterize the major axes of shape variation. Assessment of correlations between the shape distribution along PCs and the diet or sex was evaluated using ANOVA (p<0.05). Overall effects of diet and sex on shape were assessed with MANOVA, using the first PCs explaining approximately 80% of total variation for each structure. Canonical variate analysis (CVA) was used to assess group-level shape differences and generate canonical variate plots. Wireframes for illustration of the shape differences along the PC’s that correlated significantly with diet were constructed. All statistical and GM analyses were performed using the Morpho package in R (V 2022.02).

### Dense correspondence analysis

The DeCA module in 3D Slicer 5.2.2, provides a computational framework to quantify shape differences using dense, point-to-point correspondence, across surfaces. It combines homologous landmarks with dense surface registration to generate high-resolution maps of morphological variation (29,38).

Specimens were first rigidly aligned to a control specimen using manually placed landmarks, standardizing position and orientation across all meshes. The mean control shape was then computed and used as the base model for all comparisons. DeCA applies a closest-point deformation (CPD) algorithm to establish node-level correspondence between each specimen’s surface and the base model, generating a dense field of vectors that represent local shape differences at every surface point. These steps were performed separately for the mandible, cranial skeleton, and each individual muscle analyzed in this study. Outputs were visualized as heatmaps depicting the mean shape differences for the control and PLX5622 specimen. Because left–right patterns were highly similar, only right-side muscle results are presented for clarity.

## REFERENCES

1. Yeung YG, Jubinsky PT, Sengupta A, Yeung DC, Stanley ER. Purification of the colony-stimulating factor 1 receptor and demonstration of its tyrosine kinase activity. Proc Natl Acad Sci USA. 1987 Mar;84(5):1268–71.

2. Stanley ER, Chitu V. CSF-1 Receptor Signaling in Myeloid Cells. Cold Spring Harbor Perspectives in Biology. 2014 June 1;6(6):a021857–a021857.

3. Chitu V, Gokhan Ş, Nandi S, Mehler MF, Stanley ER. Emerging Roles for CSF-1 Receptor and its Ligands in the Nervous System. Trends in Neurosciences. 2016 June;39(6):378–93.

4. Stewart TA, Hughes K, Hume DA, Davis FM. Developmental Stage-Specific Distribution of Macrophages in Mouse Mammary Gland. Front Cell Dev Biol. 2019 Oct 24;7:250.

5. Banaei-Bouchareb L, Gouon-Evans V, Samara-Boustani D, Castellotti MC, Czernichow P, Pollard JW, et al. Insulin cell mass is altered in *Csf1 op /Csf1 op* macrophage-deficient mice. Journal of Leukocyte Biology. 2004 June 3;76(2):359–67.

6. Araki M, Fukumatsu Y, Katabuchi H, Shultz LD, Takahashi K, Okamura H. Follicular Development and Ovulation in Macrophage Colony-Stimulating Factor-Deficient Mice Homozygous for the Osteopetrosis (Op) Mutation1. Biology of Reproduction. 1996 Feb 1;54(2):478–84.

7. Kim JM, Lin C, Stavre Z, Greenblatt MB, Shim JH. Osteoblast-Osteoclast Communication and Bone Homeostasis. Cells. 2020 Sept 10;9(9):2073.

8. Abboud SL, Woodruff K, Liu C, Shen V, Ghosh-Choudhury N. Rescue of the Osteopetrotic Defect in *op/op* Mice by Osteoblast-Specific Targeting of Soluble Colony-Stimulating Factor-1. Endocrinology. 2002 May;143(5):1942–9.

9. Keshvari S, Caruso M, Teakle N, Batoon L, Sehgal A, Patkar OL, et al. CSF1R-dependent macrophages control postnatal somatic growth and organ maturation. Barsh GS, editor. PLoS Genet. 2021 June 3;17(6):e1009605.

10. Borycki AG, Smadja F, Stanley R, Leibovitch SA. Colony-Stimulating Factor 1 (CSF-1)Is Involved in an Autocrine Growth Control of Rat Myogenic Cells. Experimental Cell Research. 1995 May;218(1):213–22.

11. Marks SC, Lane PW. Osteopetrosis, a New Recessive Skeletal Mutation on Chromosome 12 of the Mouse. Journal of Heredity. 1976 Jan;67(1):11–8.

12. Wiktor-Jedrzejczak W, Bartocci A, Ferrante AW, Ahmed-Ansari A, Sell KW, Pollard JW, et al. Total absence of colony-stimulating factor 1 in the macrophage-deficient osteopetrotic (op/op) mouse. Proc Natl Acad Sci USA. 1990 June;87(12):4828–32.

13. Begg SK, Radley JM, Pollard JW, Chisholm OT, Stanley ER, Bertoncello I. Delayed hematopoietic development in osteopetrotic (op/op) mice. The Journal of experimental medicine. 1993 Jan 1;177(1):237–42.

14. Dai XM, Ryan GR, Hapel AJ, Dominguez MG, Russell RG, Kapp S, et al. Targeted disruption of the mouse colony-stimulating factor 1 receptor gene results in osteopetrosis, mononuclear phagocyte deficiency, increased primitive progenitor cell frequencies, and reproductive defects. Blood. 2002 Jan 1;99(1):111–20.

15. Dai XM, Zong XH, Akhter MP, Stanley ER. Osteoclast Deficiency Results in Disorganized Matrix, Reduced Mineralization, and Abnormal Osteoblast Behavior in Developing Bone. Journal of Bone and Mineral Research. 2004 Sept 1;19(9):1441–51.

16. Van Wesenbeeck L, Odgren PR, MacKay CA, D’Angelo M, Safadi FF, Popoff SN, et al. The osteopetrotic mutation *toothless* (*tl*) is a loss-of-function frameshift mutation in the rat *Csf1* gene: Evidence of a crucial role for CSF-1 in osteoclastogenesis and endochondral ossification. Proc Natl Acad Sci USA. 2002 Oct 29;99(22):14303–8.

17. Pridans C, Raper A, Davis GM, Alves J, Sauter KA, Lefevre L, et al. Pleiotropic Impacts of Macrophage and Microglial Deficiency on Development in Rats with Targeted Mutation of the *Csf1r* Locus. The Journal of Immunology. 2018 Nov 1;201(9):2683–99.

18. Dumont NA, Frenette J. Macrophage Colony-Stimulating Factor–Induced Macrophage Differentiation Promotes Regrowth in Atrophied Skeletal Muscles and C2C12 Myotubes. The American Journal of Pathology. 2013 Feb;182(2):505–15.

19. Wang X, Sathe AA, Smith GR, Ruf-Zamojski F, Nair V, Lavine KJ, et al. Heterogeneous origins and functions of mouse skeletal muscle-resident macrophages. Proc Natl Acad Sci USA. 2020 Aug 25;117(34):20729–40.

20. Hume DA, Caruso M, Ferrari-Cestari M, Summers KM, Pridans C, Irvine KM. Phenotypic impacts of CSF1R deficiencies in humans and model organisms. Journal of Leukocyte Biology. 2020 Feb 1;107(2):205–19.

21. Rosin JM, Vora SR, Kurrasch DM. Depletion of embryonic microglia using the CSF1R inhibitor PLX5622 has adverse sex-specific effects on mice, including accelerated weight gain, hyperactivity and anxiolytic-like behaviour. Brain, Behavior, and Immunity. 2018 Oct;73:682–97.

22. Spiteri AG, Ni D, Ling ZL, Macia L, Campbell IL, Hofer MJ, et al. PLX5622 Reduces Disease Severity in Lethal CNS Infection by Off-Target Inhibition of Peripheral Inflammatory Monocyte Production. Front Immunol. 2022 Mar 25;13:851556.

23. Nagra A, Katsube M, Gao W, Rosin JM, Vora SR. Embryonic inhibition of colony-stimulating factor 1 receptor impacts craniofacial morphogenesis. Orthod Craniofacial Res. 2023 Dec;26(S1):20–8.

24. Guo L, Ikegawa S. From HDLS to BANDDOS: fast-expanding phenotypic spectrum of disorders caused by mutations in CSF1R. J Hum Genet. 2021 Dec;66(12):1139–44.

25. Dulski J, Souza J, Santos ML, Wszolek ZK. Brain abnormalities, neurodegeneration, and dysosteosclerosis (BANDDOS): new cases, systematic literature review, and associations with CSF1R-ALSP. Orphanet J Rare Dis. 2023 June 22;18(1):160.

26. Daghagh H, Rahbar Kafshboran H, Daneshmandpour Y, Nasiri Aghdam M, Talebian S, Nouri Nojadeh J, et al. Homozygous mutation in *CSF1R* causes brain abnormalities, neurodegeneration, and dysosteosclerosis (BANDDOS). Bioimpacts. 2022 Nov 26;1.

27. Oosterhof N, Chang IJ, Karimiani EG, Kuil LE, Jensen DM, Daza R, et al. Homozygous Mutations in CSF1R Cause a Pediatric-Onset Leukoencephalopathy and Can Result in Congenital Absence of Microglia. The American Journal of Human Genetics. 2019 May;104(5):936–47.

28. Ma F, Zhou RRJ, Rosin M, Zhou I, Ownsworth S, Memar RO, et al. CSF1R+ macrophage and osteoclast depletion impairs neural crest proliferation and craniofacial morphogenesis [Internet]. 2025 [cited 2025 Nov 6]. Available from: http://biorxiv.org/lookup/doi/10.1101/2025.10.08.681206

29. Rolfe SM, Maga AM. DeCA: A Dense Correspondence Analysis Toolkit for Shape Analysis. In: Wachinger C, Paniagua B, Elhabian S, Li J, Egger J, editors. Shape in Medical Imaging [Internet]. Cham: Springer Nature Switzerland; 2023 [cited 2025 Jan 31]. p. 259–70. (Lecture Notes in Computer Science; vol. 14350). Available from: https://link.springer.com/10.1007/978-3-031-46914-5_21

30. Rosin JM, Sinha S, Biernaskie J, Kurrasch DM. A subpopulation of embryonic microglia respond to maternal stress and influence nearby neural progenitors. Developmental Cell. 2021 May;56(9):1326–1345.e6.

31. Rosin JM, Vora SR, Kurrasch DM. Depletion of embryonic microglia using the CSF1R inhibitor PLX5622 has adverse sex-specific effects on mice, including accelerated weight gain, hyperactivity and anxiolytic-like behaviour. Brain, Behavior, and Immunity. 2018 Oct;73:682–97.

32. Sui H, Dou J, Shi B, Cheng X. The reciprocity of skeletal muscle and bone: an evolving view from mechanical coupling, secretory crosstalk to stem cell exchange. Front Physiol. 2024 Mar 4;15:1349253.

33. Erblich B, Zhu L, Etgen AM, Dobrenis K, Pollard JW. Absence of Colony Stimulation Factor-1 Receptor Results in Loss of Microglia, Disrupted Brain Development and Olfactory Deficits. Meisel A, editor. PLoS ONE. 2011 Oct 27;6(10):e26317.

34. Tunster SJ, Van De Pette M, John RM. Fetal overgrowth in the *Cdkn1c* mouse model of Beckwith-Wiedemann syndrome. Disease Models & Mechanisms. 2011 Nov 1;4(6):814–21.

35. Metscher BD. MicroCT for developmental biology: A versatile tool for high-contrast 3D imaging at histological resolutions. Developmental Dynamics. 2009 Mar;238(3):632–40.

36. Baverstock H, Jeffery NS, Cobb SN. The morphology of the mouse masticatory musculature. Journal of Anatomy. 2013 July;223(1):46–60.

37. Vora SR, Camci ED, Cox TC. Postnatal Ontogeny of the Cranial Base and Craniofacial Skeleton in Male C57BL/6J Mice: A Reference Standard for Quantitative Analysis. Front Physiol [Internet]. 2016 Jan 12 [cited 2024 Nov 14];6. Available from: http://journal.frontiersin.org/Article/10.3389/fphys.2015.00417/abstract

38. Katsube M, Rolfe SM, Bortolussi SR, Yamaguchi Y, Richman JM, Yamada S, et al. Analysis of facial skeletal asymmetry during foetal development using μCT imaging. Orthod Craniofacial Res. 2019 May;22(S1):199–206.

