## Supplemental Figures 1 and 2 for "Gestational inhibition of CSF1R signaling using PLX5622 drives musculoskeletal changes in postnatal offspring"

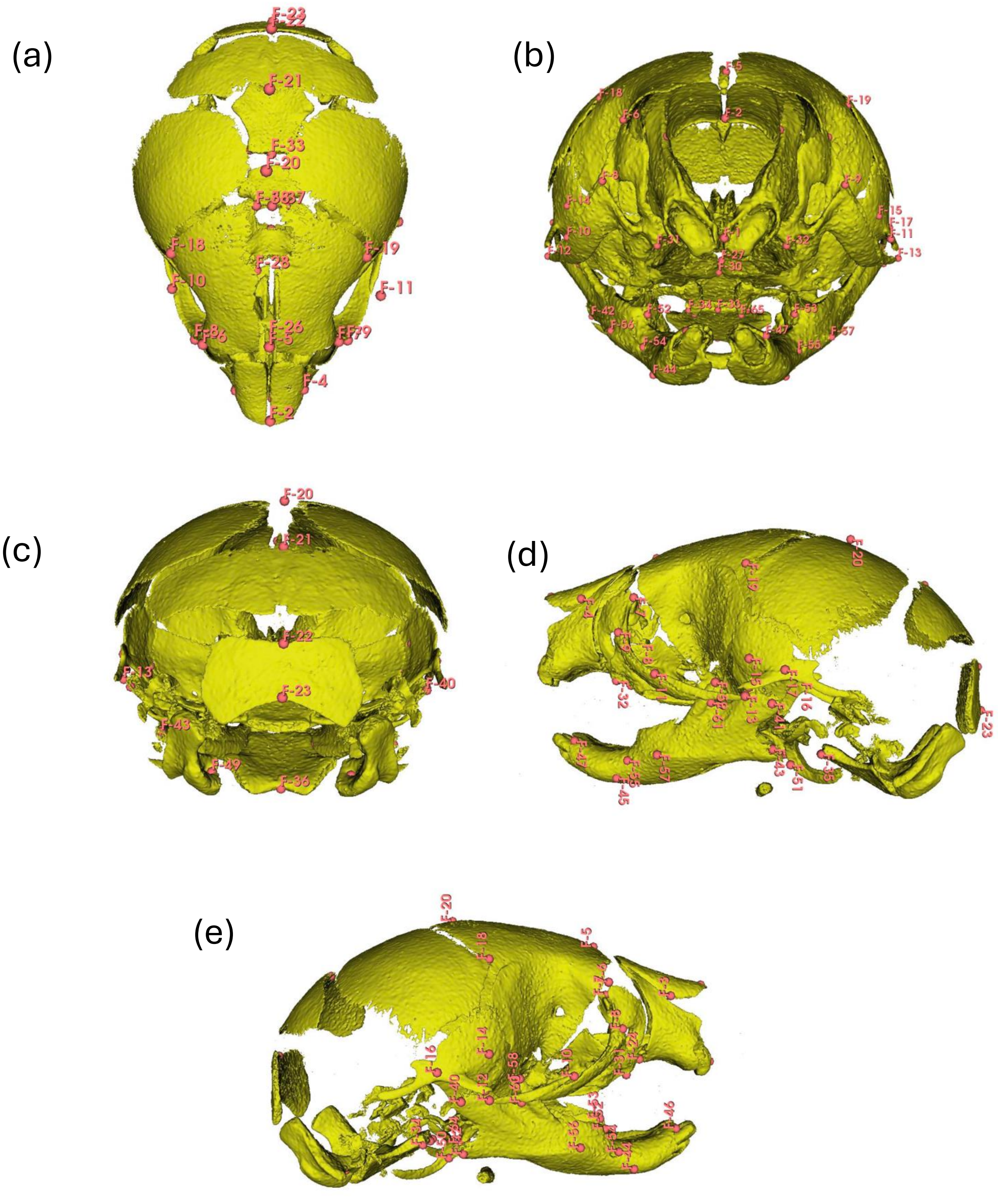
(a)
(b)
(c)
(d)
(e)

**Supplemental Figure 1. Annotated landmarks of the skull.** Superior view (A), anterior view (B), posterior view (C), left lateral view (D), and right lateral view (E) of the skull. Description for each annotated landmark is provided in Table 2.

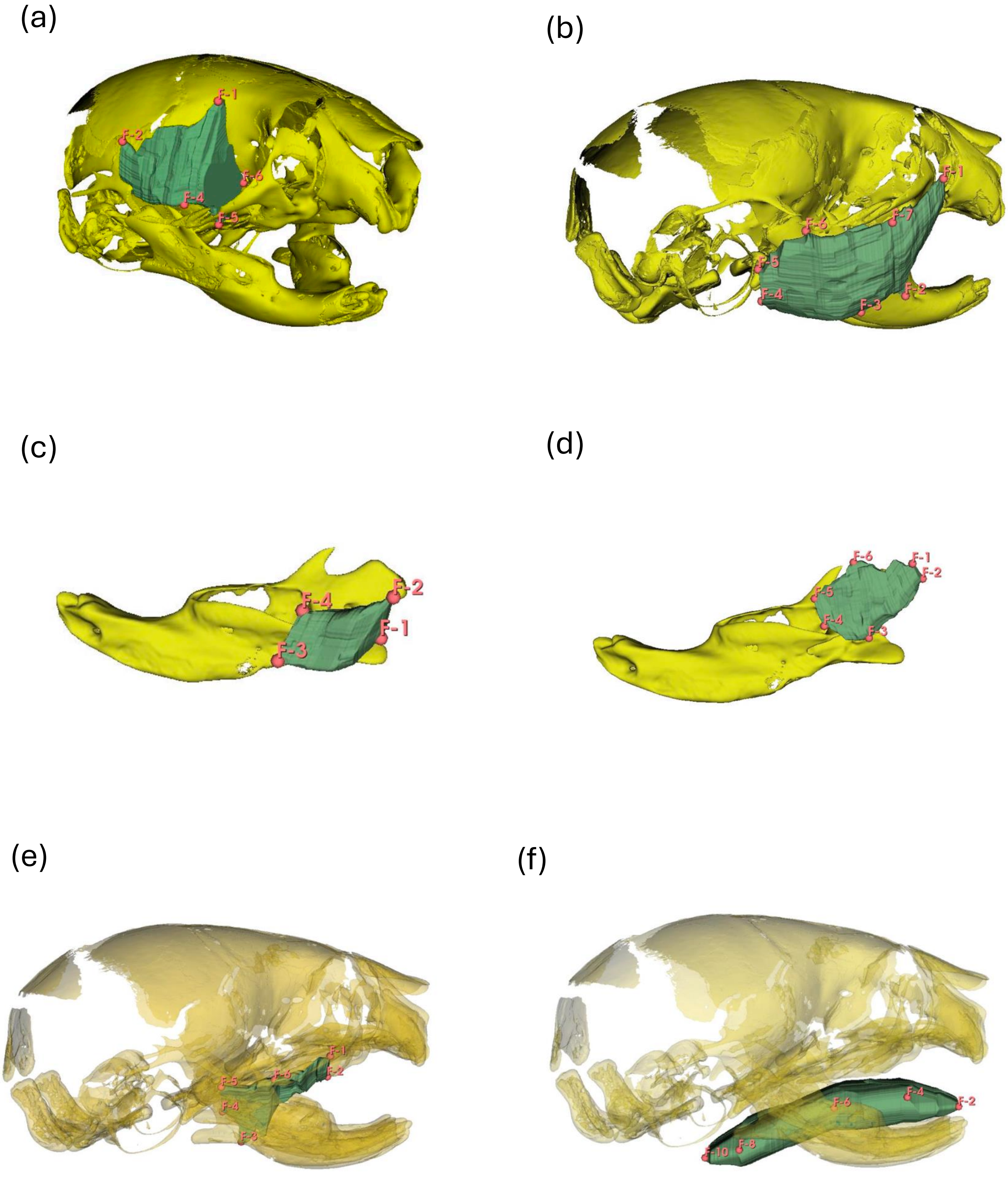

(a)
F-1
F-2
F-6
F-4
F-5
(b)
F-1
F-6
F-7
F-5
F-4
F-3
F-2
(c)
F-2
F-4
F-1
F-3
(d)
F-6
F-1
F-2
F-5
F-4
F-3
(e)
F-1
F-5
F-6
F-2
F-4
F-3
(f)
F-6
F-4
F-2
F-10
F-8

**Supplemental Figure 2. Annotated landmarks of the muscles of mastication and tongue.** (A) Right temporalis. (B) Right masseter. (C) Right internal pterygoid. (D) Right external pterygoid. (E) Right zygomatico-mandibularis. (F) Right lateral view of the tongue. Description for each annotated landmark is provided in Table 3.
